# Fine-tuning of ABA responses by the protein kinase WNK8

**DOI:** 10.1101/251975

**Authors:** Rainer Waadt, Kenji Hashimoto, Esther Jawurek, Melanie Krebs, Martin Scholz, Anne Hong-Hermesdorf, Yan Li, Michael Hippler, Erwin Grill, Jörg Kudla, Karin Schumacher

**Affiliations:** Heidelberg University, Centre for Organismal Studies, Dep. Cell Biology, Im Neuenheimer Feld 230, 69120 Heidelberg, Germany.; Westfälische-Wilhelms-Universität Münster, Institut für Biologie und Biotechnologie der Pflanzen, Schlossplatz 7, 48149, Münster, Germany.; Westfälische-Wilhelms-Universität Münster, Institut für Biologie und Biotechnologie der Pflanzen, Schlossplatz 8, 48143 Münster, Germany; Technische Universität München, Lehrstuhl für Botanik, Emil-Ramann-Straße 4, 85354 Freising, Germany.

**Author notes:** Present address: Tokyo University of Science, Department of Applied Biological Science, 2641 Yamazaki, Noda 278-8510, Japan (Kenji Hashimoto). Present address: University of California, Los Angeles, Department of Chemistry and Biochemistry, 607 Charles E. Young Drive East, Los Angeles, CA 90095-1569, USA. **CORRESPONDING AUTHOR:** Karin Schumacher Heidelberg University Centre for Organismal Studies Dep. Cell Biology Im Neuenheimer Feld 230 69120 Heidelberg Germany. **email of all authors:**.

**Keywords:** Abscisic Acid, Arabidopsis thaliana, protein kinase WNK8, protein phosphatase PP2CA

## Abstract

The phytohormone abscisic acid (ABA) regulates various growth- and developmental processes in response to limiting water conditions. ABA functions through an established signaling pathway consisting of PYR1/PYL/RCAR ABA receptors that inhibit group A type 2C protein phosphatases (PP2Cs) in an ABA-dependent manner. Inhibition of PP2Cs enables the activation of SnRK2-type protein kinases that phosphorylate downstream targets including transcription factors and ion channels. However, ABA-dependent signals have to be integrated into other growth- and developmental programs to ensure a successful life cycle. Here, we have characterized the role of the protein kinase WNK8 in the ABA signalling pathway. Two T-DNA insertion alleles *wnk8-1* and *wnk8-4* exhibited contrasting ABA responses during seed germination and young seedling growth. However, reciprocal crossings with wild type plants suggested that *wnk8-1* that still expressed the WNK8 kinase domain functioned in a hypermorphic and dominant manner. WNK8 directly interacted with the PP2C PP2CA *in planta* and was negatively regulated by this phosphatase *in vitro*. WNK8 also phosphorylated the ABA receptor PYR1 *in vitro*. Double mutant analyses revealed that the dominant allele *wnk8-1* suppressed the ABA- and glucose hypersensitivity of the *pp2ca-1* T-DNA allele. In transient protoplast assays WNK8 suppressed ABA-induced reporter gene expression that was dependent on a functional kinase. In summary, we have identified the protein kinase WNK8 as a negative regulator of ABA responses during young seedling establishment through its direct interaction with core ABA signaling components.

**SIGNIFICANCE STATEMENT:** The phytohormone abscisic acid regulates the water household of plants through a defined core signaling pathway. Here we have identified the protein kinase WNK8 as a direct interactor of core abscisic acid signalling components and as a negative modulator of abscisic acid responses during young seedling development in Arabidopsis.

## INTRODUCTION

Phosphorylation by protein kinases is the most widespread and well-studied signaling mechanism and all eukaryotic genomes encode large numbers of protein kinases that fall into many divergent subclasses but share the strictly organized internal architecture of their kinase domains (Taylor & Kornev, 2011). WNK (with no lysine (K)) kinases form an ancient clade of the eukaryotic kinome characterized by the unusual position of their catalytic lysine residue in strand 2 (subdomain I) instead of the canonical position in strand 3 (subdomain II) found in all other kinase families (Xu et al., 2000; Min et al., 2004). Outside of the kinase domain the only conserved feature found in most members of the WNK family is a short autoinhibitory domain (AID) embedded in a C-terminal extension of highly variable size and sequence (Hong-Hermesdorf et al., 2006). The Arabidopsis genome encodes for 11 members of the WNK-family (Hong-Hermesdorf et al., 2006). Based on loss-of-function mutants, WNKs appeared to be involved in the regulation of flowering time (Wang et al., 2008) and in glucose-, drought- and ABA responses (Urano et al., 2012; Zhang et al., 2013; Fu et al., 2014; Xie et al., 2014).

A number of proteins have been identified as interaction partners and phosphorylation substrates of WNK8, making it the so far best studied family member. WNK8 was reported to bind and phosphorylate VHA-C, a subunit of the vacuolar H^+^-ATPase (Hong-Hermesdorf et al., 2006), the nitrate transceptor NRT1.1 (Jones et al., 2014; Ho et al., 2014), the transcriptional regulator EDM2 (Tsuchiya and Eulgem, 2010), the G protein-coupled receptor RGS1 (Urano et al., 2012), and the scaffold protein RACK1 (Urano et al., 2015). Among these, RACK1 genes have been implicated as a negative regulators of ABA responses (Guo et al., 2009; 2011). However, a molecular link between ABA-signaling components and RACK1 has not been established.

ABA signals are perceived by ABA-binding Pyrabactin Resistance 1/Pyrabactin Resistance 1-Like/Regulatory Component of ABA Receptor (PYR1/PYL/RCAR) proteins that interact with and, in the presence of ABA, inhibit group A Type 2C Protein Phosphatases (PP2Cs) that function as ABA co-receptors (Ma et al., 2009; Park et al., 2009). Group A PP2Cs are negative regulators of ABA signaling that de-phosphorylate and inhibit Sucrose Non-Fermenting 1 (SNF1) Related Protein Kinases of the subgroup 2 (SnRK2s) (Umezawa et al., 2009; Vlad et al., 2009). PYR1/PYL/RCAR and ABA-mediated inhibition of PP2Cs enables the re-activation of SnRK2s that phosphorylate downstream targets to regulate seed dormancy and germination, stomatal movements and gas exchange and root growth and development (Cutler et al., 2010; Raghavendra et al., 2010; Hubbard et al., 2010). This core ABA signaling pathway is further interconnected with other signaling pathways through direct protein-protein interactions, protein phosphorylation/de-phosphorylation events or transcriptional responses (Lumba et al., 2014; Umezawa et al., 2013; Wang et al., 2013; Waadt et al., 2015).

PP2Cs can interact with and de-phosphorylate several proteins, including direct substrates of SnRK2s (reviewed in Edel and Kudla, 2016). However, group A PP2Cs also modulate the activity of other protein kinases. For example, SnRK1.1, a member of the subgroup 1 of SnRK-type protein kinases, that is involved in sugar responses, interacts with and is de-phosphorylated by the group A PP2Cs ABI1 and PP2CA (Rodrigues et al., 2013). Group A PP2Cs also interact with subgroup 3 SnRK-type protein kinases, designated as CBL-Interacting Protein Kinases (CIPKs) (Guo et al., 2002; Ohta et al., 2003) and with their regulating Calcineurin B-Like (CBL) calcium sensor proteins (Lan et al., 2011; Léran et al., 2015). Here, the group A PP2C ABI2 interacted with and de-phosphorylated the CBL1/CIPK23 module (Léran et al., 2015). Recently, it has been reported that the group A PP2C ABI2 directly interacted with and de-phosphorylated the kinase domain of the Receptor-Like Kinase FERONIA (FER) (Chen et al., 2016).

PP2CA is a member of the Arabidopsis core ABA signalling network group A PP2Cs (Schweighofer et al., 2004). PP2CA is located in the nucleus (Umezawa et al., 2009), and it interacts with and de-phosphorylates SnRK2s (Park et al., 2009; Lee et al., 2009; Brandt et al., 2015) as well as their substrate transcription factors, the ABA-Responsive Element (ABRE)-Binding Proteins (AREB)/ABRE-Binding Factors (ABFs) (Lynch et al., 2012) and the SWI/SNF chromatin-remodeling ATPase BRAHMA (Peirats-Llobet et al., 2015). Although located in the nucleus (Umezawa et al., 2009), there are also reports from *in vitro* or heterologous analyses in which PP2CA targeted the K^+^ channels AKT1, AKT2 and GORK (Cherel et al., 2002; Lan et al., 2011; Lefoulon et al., 2016) and the anion channel SLAC1 (Lee et al., 2009; Brandt et al., 2015). PP2CA is involved in all classical ABA-mediated responses, including inhibition of seed germination, root growth and stomatal closure, with loss-of-function alleles displaying ABA hypersensitive phenotypes (Kuhn et al., 2006; Yoshida et al., 2006). Among the group A PP2Cs, PP2CA plays a prominent role during seed germination and it was identified in a forward genetic screen for ABA Hypersensitive Germination (AHG3) (Nishimura et al., 2004; Yoshida et al., 2006).

Here we identified the protein kinase WNK8 as a negative regulator of ABA responses. WNK8 physically interacted with PP2CA in Bimolecular Fluorescence Complementation (BiFC) and Förster Resonance Energy Transfer (FRET)-Fluorescence Lifetime (FLIM) assays *in planta* and was negatively regulated by PP2CA *in vitro*. Apart from that, WNK8 phosphorylated the ABA receptor PYR1 *in vitro*, but not PYL8. Arabidopsis protoplast ABA response assays revealed that WNK8 functions as a negative regulator of ABA signalling. This was also supported by seed germination and young seedling growth analyses of WNK8 T-DNA insertion alleles. Although two *wnk8* T-DNA alleles *wnk8-1* and *wnk8-4* exhibited opposite ABA responses, reciprocal crossing with Col-0 wild type suggested that the *wnk8-1* allele, harbouring the T-DNA insertion in the kinase regulatory domain, behaved as a dominant hypermorphic allele.

## RESULTS

### WNK8 is expressed during young seedling development and is involved in ABA-mediated responses during that developmental stage

Gene expression analyses using the Arabidopsis eFP browser (Winter et al., 2007) and Genevestigator (Hruz et al., 2008) indicated that the *WNK8* gene (AT5G41990) is highly expressed in pollen, chalazal tissues and the root vasculature. *Promoter::GUS* analyses using a promoter fragment of−1868 bp to +9 bp from the *WNK8* start codon indicated *WNK8* expression in two-day-old seedlings (Figure S1a), in roots, root and shoot apices, the hypocotyl and cotyledon vasculature of five-day-old seedlings (Figure S1b), in guard cells of cotyledons (Figure S1c) and in the sepal vasculature, anther filaments and the flower stigma (Figure S1d). The observed expression pattern is reminiscent of genes involved in ABA synthesis, transport and signaling (Boursiac et al., 2013) and we thus asked whether *WNK8* is involved in ABA responses. To test this hypothesis, we characterised two T-DNA insertion alleles, *wnk8-1* (SALK_024887; Wang et al., 2008, *wnk8-2* in Urano et al., 2012) with insertion in the sequence encoding for the C-terminal regulatory region and *wnk8-4* (SALK_103318 in Wang et al., 2008; *wnk8-1* in Urano et al., 2012) with insertion in the kinase domain (Figure 1a). Both T-DNA alleles did not express full-length *WNK8* transcripts. However, the *wnk8-1* allele still expressed the kinase domain and the *wnk8-4* allele expressed transcripts 3’ of the insertion site (Figure 1b).

**Figure 1.**
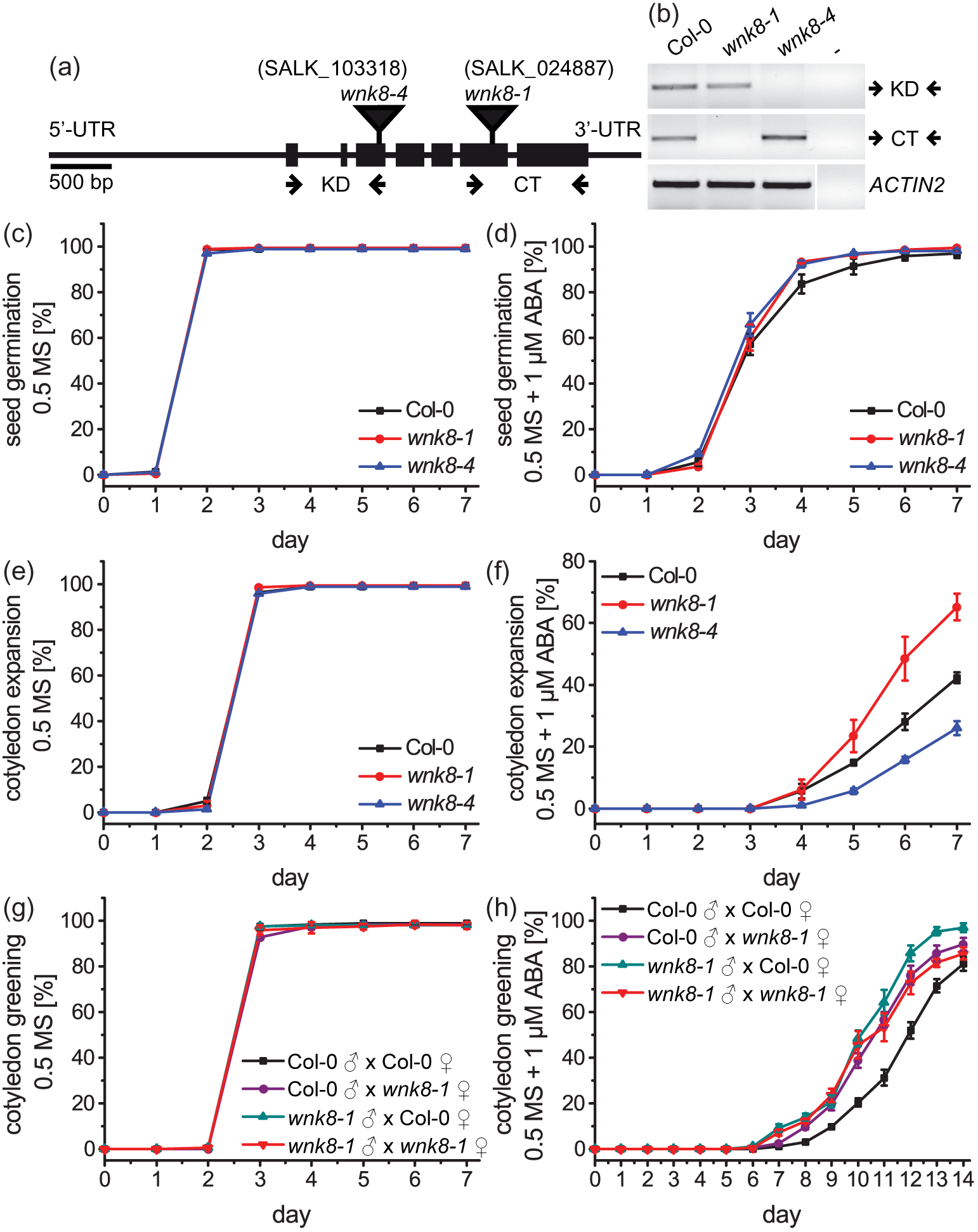
Mutations in the *WNK8* gene affect ABA responses during young seedling development. (a) Schematic presentation of the *WNK8* gene. 5’- and 3’-untranslated regions (UTR) and introns are depicted by lines and exons are indicated by boxes. T-DNA insertion sites of respective *wnk8* alleles are indicated by inverted triangles. Positions of oligo nucleotides used for RT-PCRs in (b) for the amplification of the *WNK8* kinase domain (KD) or C-terminal region (CT) are indicated by arrows. (b) RT-PCRs of Col-0 wild type, indicated *wnk8* alleles and buffer control (-). PCRs were performed for the amplification of the WNK8 kinase domain (KD) and C-terminal region (CT; indicated in (a)). *ACTIN2* was used as a positive control. (c,d) Time course of seed germination, (e,f) cotyledon expansion or (g,h) cotyledon greening of indicated genotypes when grown on 0.5 MS, 1% sucrose media supplemented with (c,e,g) 0.1% EtOH as solvent control or (d,f,h) 1 μM ABA. Data indicate means ± SEM of n = 4 experiments.

Using seeds of parallel grown and similarly treated wild type Col-0 and *wnk8* alleles we initially investigated the kinetics of ABA-dependent seed germination (Figure 1c,d) and cotyledon expansion (Figure 1e,f) when grown on control media (0.5 MS + 1% sucrose + 0.1% EtOH) or control media supplemented with 1 μM ABA dissolved in EtOH. No altered seed germination was observed. However, in response to 1 μM ABA, cotyledon expansion of *wnk8-1* appeared faster compared to Col-0 wild type, while *wnk8-4* cotyledons expanded slower (Figure 1f).

The fact that *wnk8-1* and *wnk8-4* exhibited opposite phenotypes was reminiscent to other similar findings about the Receptor-Like Kinase (RLK) THESEUS1 (Merz et al., 2017) and implied that only one of the two *wnk8* alleles might be a real loss-of-function allele. As transcripts encoding the WNK8 kinase domain but not the autoinhibitory domain were detectable in *wnk8-1*, we reasoned that this allele could function in a dominant fashion through the expression of a constitutively active WNK8 kinase. To test this hypothesis we crossed Col-0 wild type and *wnk8-1* plants in all possible orientations and subjected F1 seeds to a more stringent assay, where we counted seedlings with expanded and green cotyledons (Figure 1g,h). In these assays, F1 progenies that harbour one or both *wnk8-1* alleles exhibited a reduced ABA sensitivity compared to Col-0 wild type, indicating a dominant nature of the *wnk8-1* allele (Figure 1h).

### WNK8 interacts with the group A type 2C protein phosphatase PP2CA in the nucleus

Due to the role of WNK8 during young seedling establishment in response to ABA, we suspected that WNK8 might directly interact with core ABA signaling components such as the PYR1/PYL/RCAR ABA receptors or the group A PP2C co-receptors. Among these, genes encoding for the ABA receptor PYR1 and the PP2Cs ABI1, ABI2, AHG1 and AHG3/PP2CA have been isolated in genetic screens for mutants defective in ABA-mediated responses during seed germination (Koornneef et al., 1984; Meyer et al., 1994; Rodriguez et al., 1998; Nishimura et al., 2004; 2007; Yoshida et al., 2006; Park et al., 2009). Therefore, these genes appear to be the key ABA receptor and co-receptor components that function during seed germination.

To investigate whether WNK8 forms protein complexes with the PP2Cs PP2CA and ABI1 and the ABA receptors PYR1/RCAR11 and PYL8/RCAR3 and whether WNK8 forms homomers we performed *in planta* BiFC analyses (Walter et al., 2004; Waadt et al., 2008) after transient co-expression in *Nicotiana benthamiana* leaf epidermis cells (Figure 2). We also assayed the complex formation of PYR1 with PP2CA (Park et al., 2009) as positive control and combination of WNK8 with the red fluorescent protein mCherry (Shaner et al., 2004) as negative control.

**Figure 2.**
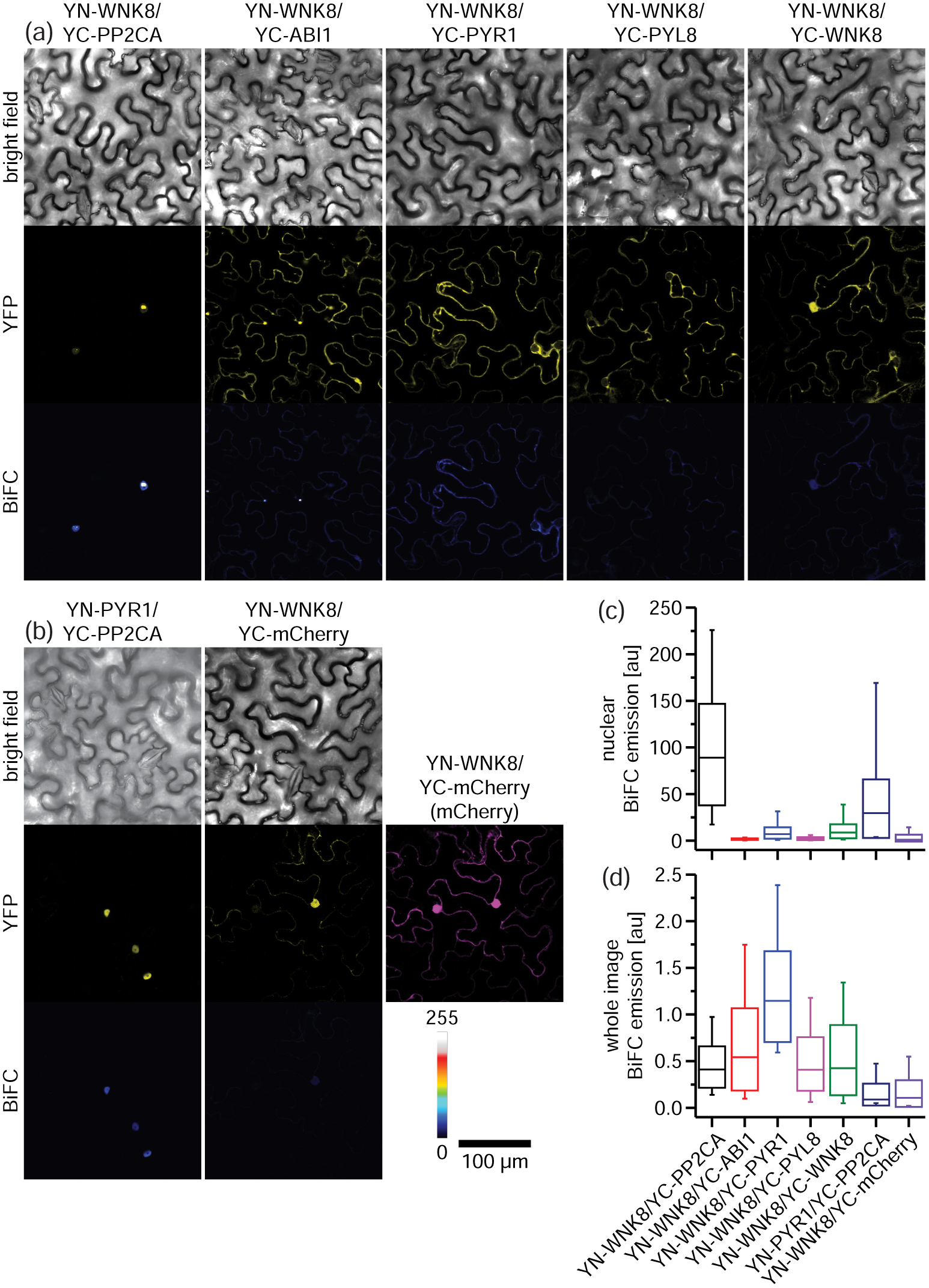
WNK8 interacts with PP2CA in BiFC analyses. (a,b) From top to bottom, representative bright field, subcellular localization (YFP) and comparative BiFC emission (BiFC) of indicated protein complex combinations after transient expression in *N. benthamiana* leaf epidermis cells. BiFC emissions were calibrated to the adjacent color bar. Expression of YC-mCherry is indicated in magenta. (c,d) Quantification of (c) nuclear- and (d) whole image BiFC emissions from indicated protein complex combinations. Boxes, median ± SD; whiskers, minimum and maximum values (n ≥ 22 nuclei or n = 20 images). See also Figure S2 for related BiFC analyses.

In BiFC analyses N- and C-terminal fragments of fluorescent proteins (here yellow fluorescent protein (YFP) fragments YN and YC) are fused to proteins of interest and physical interaction of these proteins leads to the irreversible reconstitution of the fluorescent protein. The resulting YFP fluorescence is indicative for the strength of interaction and can also be used to determine the sub-cellular localization of the protein complexes (Kerppola, 2008). Analyses of protein complex-dependent YFP fluorescence complementation indicated that all investigated protein combinations exhibited some degree of YFP fluorescence (Figure 2a,b; YFP images). Here, reconstituted YFP fluorescence was observed predominantly in the nucleus for YN-WNK8/YC-PP2CA and YN-PYR1/YC-PP2CA combinations, while WNK8 complexes with ABI1, PYR1 and PYL8 were predominantly cytoplasmic. WNK8 homomer and YN-WNK8/YC-mCherry complexes were detectable in both the cytoplasm and the nucleus. Because complex formation of BiFC fragments is irreversible, it is required to perform quantitative BiFC analyses to substantiate the observed protein-protein interactions (Kerppola, 2008; Waadt et al., 2014). We calibrated our images to the color bar depicted in Figure 2b and also measured nuclear and whole image BiFC emissions (Figure 2c,d). The data indicated that nuclear BiFC signals of YN-WNK8/YC-PP2CA complexes were even higher than the BiFC signals of the positive control YN-PYR1/YC-PP2CA, while the other protein combinations exhibited a rather low nuclear BiFC emission (Figure 2c). Whole image BiFC quantifications indicated that WNK8 might also form complexes with PYR1 in the cytoplasm (Figure 2d). PP2CA fused to green fluorescent protein (GFP) was predominantly localized to the nucleus (Umezawa et al., 2009; Figure S2a), while GFP-WNK8 and GFP-PYR1 were localized in the cytoplasm and nucleus, similar to GFP or mCherry alone (Figure S2a). Due to the nuclear localization of PP2CA, complexes with WNK8 and PYR1 may form predominantly in the nucleus (Figure 2a,b; Figure S2b). To substantiate these nuclear interactions we performed additional BiFC analyses at higher magnification and quantified nuclear BiFC emissions (Figure S2c). The data clearly indicated higher BiFC emissions of YN-WNK8/YC-PP2CA and YN-PYR1/YC-PP2CA complexes compared to the negative control YN-WNK8/YC-mCherry.

As an alternative method to proof the physical interaction between WNK8 and PP2CA, we performed FRET-FLIM analyses in nuclei of transiently transformed *Nicotiana benthamiana* leaf epidermis cells (Figure 3). In FRET-FLIM analyses, energy transfer from a donor to an acceptor fluorescent protein is quantified, which depends on the distance (2-8 nm) and orientation of the fluorophores. Because FRET occurs only at these short distances, FRET signals are considered to be indicative for specific protein-protein interactions (Padilla-Parra and Tramier, 2012). In FRET-FLIM analyses, FRET leads to a decrease in lifetime of the donor fluorophore. This method is preferred among FRET-based protein-protein interaction analyses, as it is independent of the protein concentration (Padilla-Parra and Tramier, 2012; Bücherl et al., 2014).

**Figure 3.**
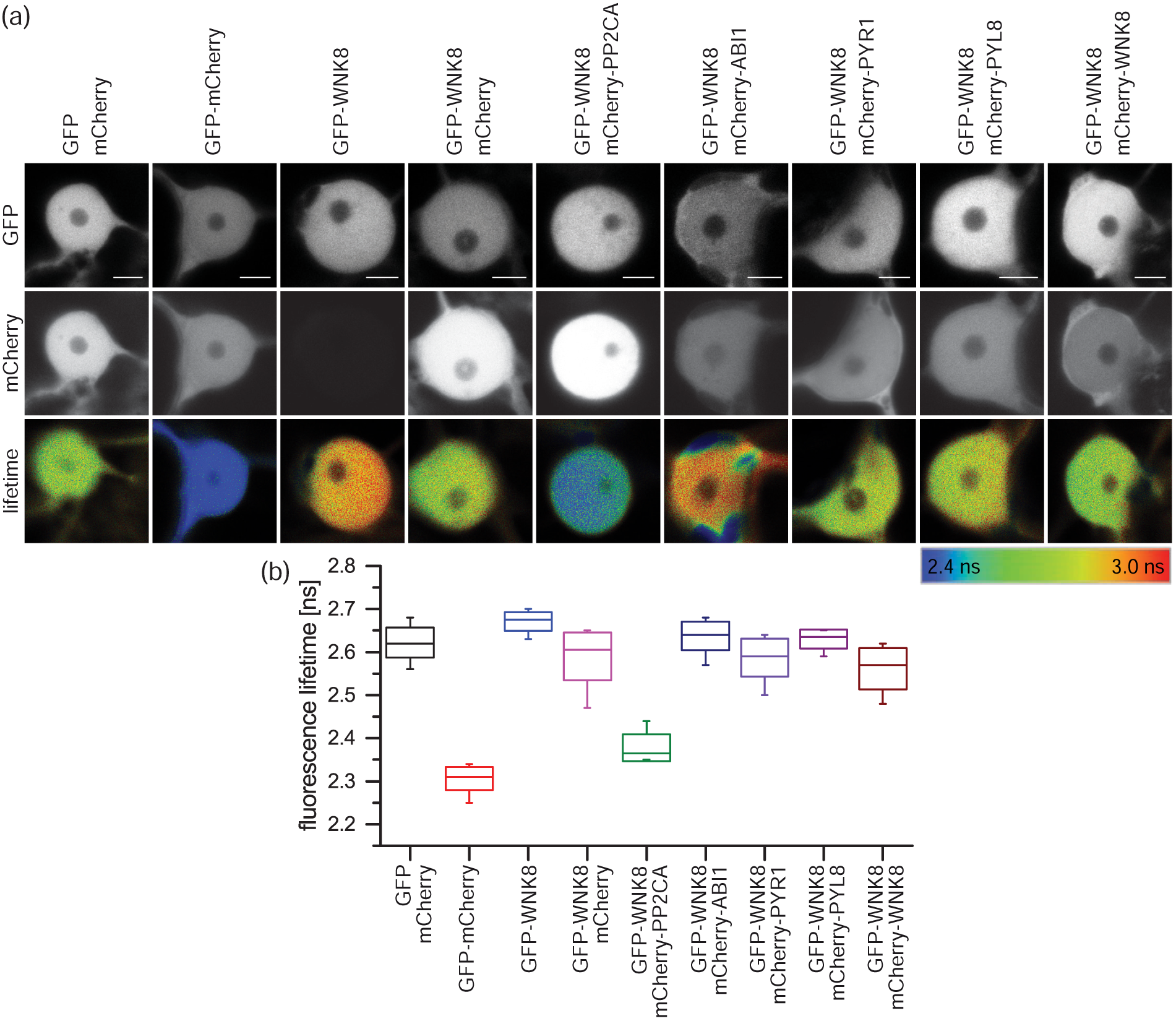
WNK8 interacts with PP2CA in the nucleus analyzed by FRET-FLIM. (a) From top to bottom, representative images of nuclear GFP, mCherry and the corresponding GFP donor fluorescence lifetime calibrated to the adjacent color bar. The respective construct combinations are indicated. Experiments were conducted after transient expression in *N. benthamiana* leaf epidermis cells. (b) GFP donor fluorescence lifetime quantifications of indicated construct combinations. Boxes, median ± SD; whiskers minimum and maximum values (n ≥ 8).

Here we used GFP as donor and mCherry as acceptor. To get better control on WNK8 expression we expressed GFP- and mCherry-WNK8 fusion proteins under control of a beta-estradiole inducible promoter (Schlücking et al., 2013). In our analyses the negative controls GFP/mCherry, GFP-WNK8 and GFP-WNK8/mCherry exhibited GFP lifetimes at about 2.59 - 2.67 ns, while the GFP-mCherry fusion, used as positive control, exhibited a GFP lifetime of 2.31 ± 0.03 ns (Figure 3a,b). GFP-WNK8 co-expressed with mCherry-PP2CA resulted in a GFP lifetime of 2.38 ± 0.03 ns, clearly decreased compared to the negative controls and indicative for a positive interaction of WNK8 with PP2CA in the nucleus (Figure 3a,b). The other construct combinations exhibited GFP lifetimes between 2.56 - 2.64 ns, comparable to the negative controls. In summary, our data strongly point towards a physical interaction between WNK8 and PP2CA in the nucleus. WNK8 might also interact with PYR1 in the cytoplasm (Figure 2a,d).

### Genetic interaction of WNK8 with PP2CA

The physical interaction of WNK8 and PP2CA prompted us to investigate whether they also interact genetically. Therefore, we crossed both *wnk8-1* and *wnk8-4* alleles with the T-DNA insertion line *pp2ca-1* (SALK_028132; Kuhn et al., 2006). Mutations in the *PP2CA/AHG3* gene render plants hypersensitive to ABA (Nishimura et al., 2004; Yoshida et al., 2006; Kuhn et al., 2006). Therefore, we modified our germination assay media and monitored seed germination and cotyledon expansion of Col-0 wild type, the single *wnk8-1*, *wnk8-2*, *pp2ca-1* and the respective double mutants on 0.5 MS media without sucrose and supplemented with 0.3 μM ABA or 0.03% EtOH as solvent control. When grown on control media, all genotypes developed in a similar manner (Figure 4a,c). Consistent with results presented in Figure 1, *wnk8-1* was less ABA sensitive than Col-0 wild type and *wnk8-4* exhibited a slightly enhanced ABA sensitivity (Figure 4b,d). Note that due to the low ABA concentration (0.3 μM) these phenotypes (Figure 4b,d) were less pronounced than the cotyledon expansion phenotype in response to 1 μM ABA (Figure 1f). Consistent with previous reports (Kuhn et al., 2006), *pp2ca-1* exhibited a greatly enhanced ABA sensitivity during seed germination and young seedling establishment (Figure 4b,d). Interestingly, the *pp2ca-1/wnk8-1* double mutant exhibited an intermediate phenotype compared to the single mutants, while growth of the *pp2ca-1/wnk8-4* double mutant was more comparable to the *pp2ca-1* single mutant (Figure 4b,d).

**Figure 4.**
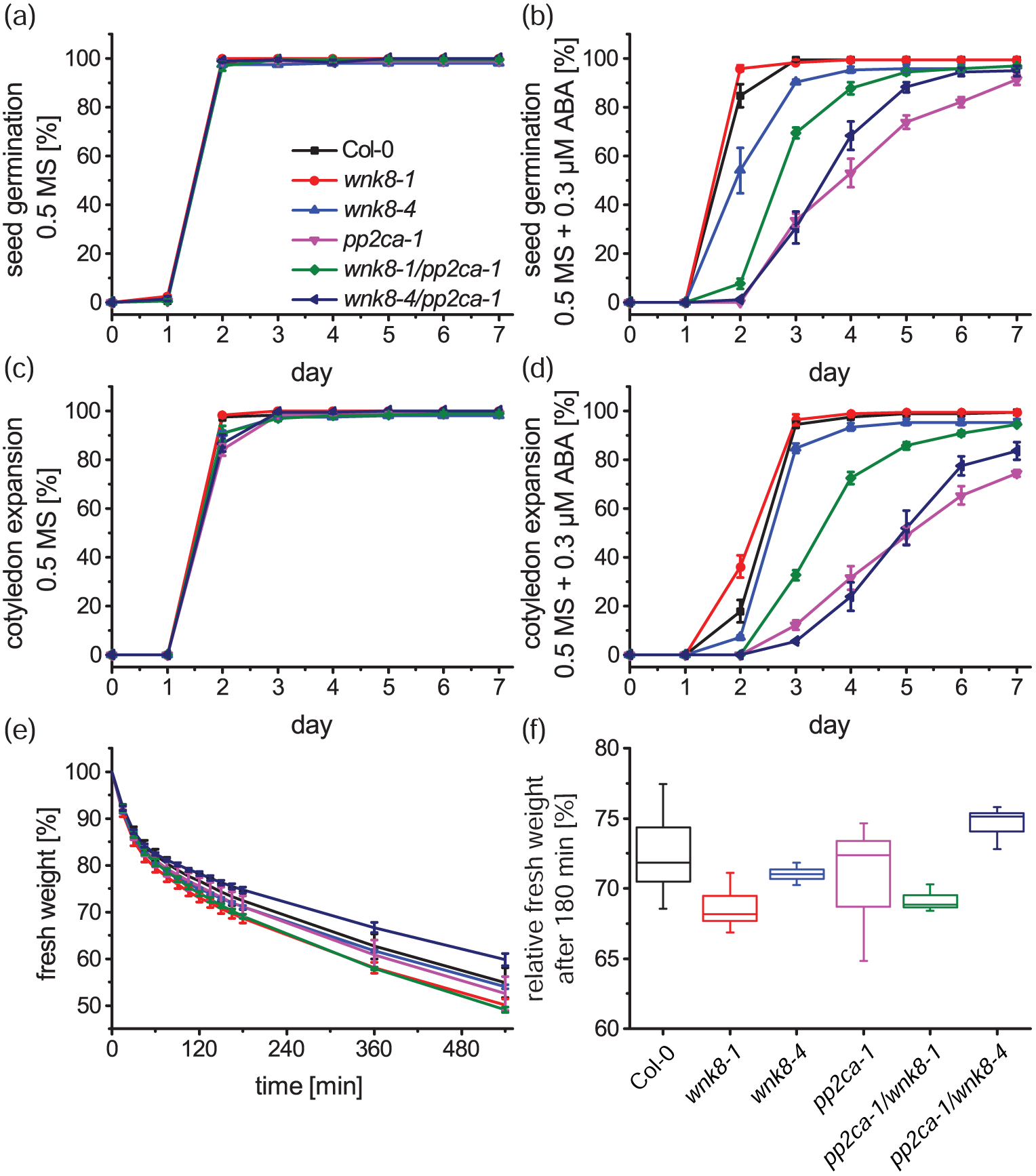
*WNK8* interacts genetically with *PP2CA*. (a,b) Time course of seed germination and (c,d) cotyledon expansion of indicated genotypes when grown on 0.5 MS media supplemented with (a,c) 0.03% EtOH as solvent control, or (b,d) 0.3 μM ABA (means ± SEM, n = 4). (e) Time course of detached rosette fresh weight of indicated genotypes normalized to the initial weight (means ± SEM, n = 4). (f) Rosette fresh weight of indicated genotypes 180 min after being detached and relative to the initial weight. Boxes, median ± SD; whiskers, minimum and maximum values (n = 4). See also Figure S3 for related experiments.

It has been previously reported that *wnk8-1* exhibits a reduced sensitivity to 6% glucose during seedling establishment (Urano et al., 2015). Based on these findings and to study the interconnection between glucose sensing and ABA we performed additional seed germination analyses on 0.5 MS media supplemented with 6% glucose and on media supplemented with 4% glucose + 150 nM ABA (Figure S3). In our experimental conditions, 6% glucose in the media had no dramatic effect on seed germination and cotyledon expansion of *wnk8-1*, while *wnk8-4* and *pp2ca-1* displayed a glucose hypersensitive phenotype with a one-day delay in growth (Figure S3b,e,g,h). Interestingly, double mutant analyses indicated that the *wnk8-1* allele in the *pp2ca-1* mutant background rescued the glucose hypersensitivity of *pp2ca-1*, but the *wnk8-4* allele did not (Figure S3b,e,g,h). In response to 4% glucose + 150 nM ABA both *wnk8* alleles exhibited opposing phenotypes, with *wnk8-1* being hyposensitive compared to Col-0 wild type and *wnk8-4* being hypersensitive (Figure S3c,f,g-i). These opposing phenotypes were consistent with data presented in Figure 1f. For the time until cotyledon expansion, *pp2ca-1* was hypersensitive to 4% glucose + 150 nM ABA and the double mutants exhibited again opposing phenotypes compared to the *pp2ca-1* background (Figure S3f,i).

PP2CA functions not only during young seedling establishment, but also in guard cells (Kuhn et al., 2006; Brandt et al., 2015). *pp2ca-1* did not directly affect fresh weight loss of detached rosettes (Figure 4e,f; Kuhn et al., 2006), most likely by water evaporation through open stomata. However, when crossed with *wnk8-1*, water loss was enhanced and when crossed with *wnk8-4*, water loss was decreased (Figure 4e,f). Note that these opposing phenotypes were less pronounced in the single *wnk8* mutant alleles.

Altogether, our phenotypic analyses indicated that *WNK8* and *PP2CA* genetically interact and that *wnk8-1* is as a hypermorphic allele of *WNK8*. The finding that in certain experimental conditions hypermorphic and dominant *wnk8-1* induced ABA hyposensitivity and *wnk8-4* induced ABA hypersensitivity support the hypothesis that *WNK8* functions as a negative regulator of ABA responses.

### WNK8 negatively affects ABA-induced gene expression

To further proof our hypothesis of WNK8 being a negative regulator of ABA signaling we performed semi-quantitative ABA-induced gene expression analyses in transiently transformed Arabidopsis protoplasts (Figure 5). Here, transfection of the ABA-responsive *pRD29B::Luciferase (LUC)* reporter into Arabidopsis protoplasts and co-expression of effector proteins, such as ABA signaling components, can be used for *in vivo* quantifications of ABA-induced gene (LUC) expression (Moes et al., 2008; Ma et al., 2009; Fujii et al., 2009). In such assays, PP2CA effectively down-regulated the ABA-induced gene expression mediated by RCAR7/PYL13 (Fuchs et al., 2014). We performed similar assays and analyzed the effect of wild type WNK8 and the inactive kinase WNK8_K41M_ (Hong-Hermesdorf et al., 2006) on PYR1 and PYL8 mediated *pRD29B::LUC* expression (Figure 5). Here, co-expression of *pRD29B::LUC* with PYR1 or PYL8 effectively induced ABA responses in the presence of 10 μM ABA. Additional co-expression of GFP-WNK8 inhibited these responses, while the kinase inactive GFP-WNK8_K41M_ did not (Figure 5). These results further supported the hypothesis that WNK8 functions as a negative regulator of ABA signaling and indicated that active WNK8 protein kinase is required for ABA response suppression.

**Figure 5.**
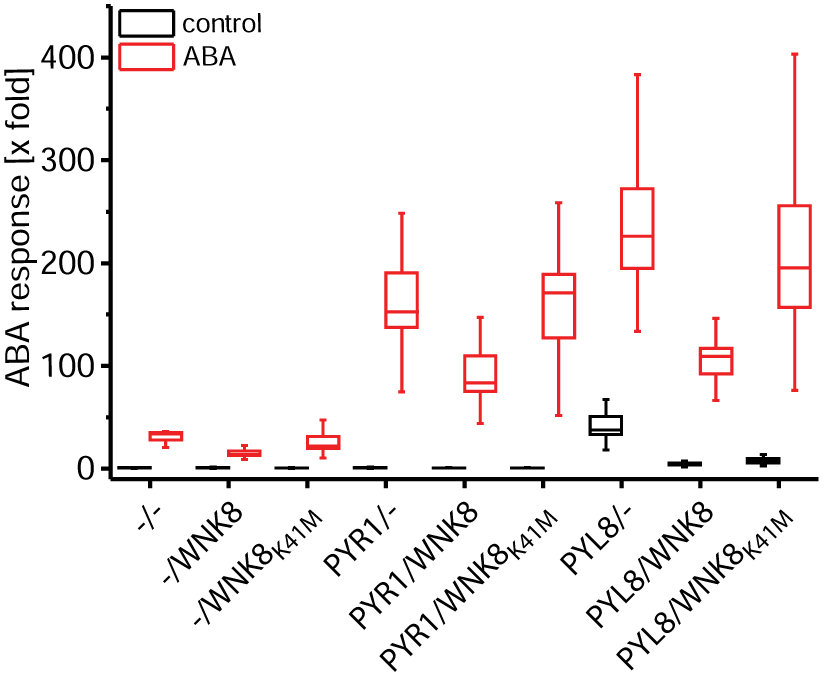
WNK8 negatively affects ABA responses in Arabidopsis protoplasts. Arabidopsis protoplasts were transfected with the *pRD29B::LUC* reporter, *p35S::GUS*, for internal normalization, and the indicated effectors. Protoplasts were incubated for 16 to 18 h without or with 10 μM ABA, and LUC luminescence was measured and normalized to the control without ABA addition. Boxes, median ± SD; whiskers, minimum and maximum values (n = 3 biological replicates).

### WNK8 phosphorylates PYR1 *in vitro* and its activity is inhibited by PP2CA

Because WNK8 interacted *in vivo* with PP2CA and potentially also with PYR1 (Figure 2, Figure 3), and negatively affected ABA responses mediated by PYR1 and PYL8 (Figure 5), we suspected that the WNK8 kinase might directly act on these core ABA signaling components. To proof whether WNK8 interacts biochemically with PP2CA, PYR1 and PYL8, we performed *in vitro* kinase assays (Figure 6). For this, StrepII-tagged WNK8 and PP2CA were expressed and purified from wheat germ extracts according to Hashimoto et al. (2012), and His-tagged PYR1 and PYL8 were expressed and purified from *E. coli* according to Ma et al. (2009) and Santiago et al. (2009b). Kinase reactions were conducted for 30 min at 30°C and reactions were started by the final addition of ATP.

**Figure 6.**
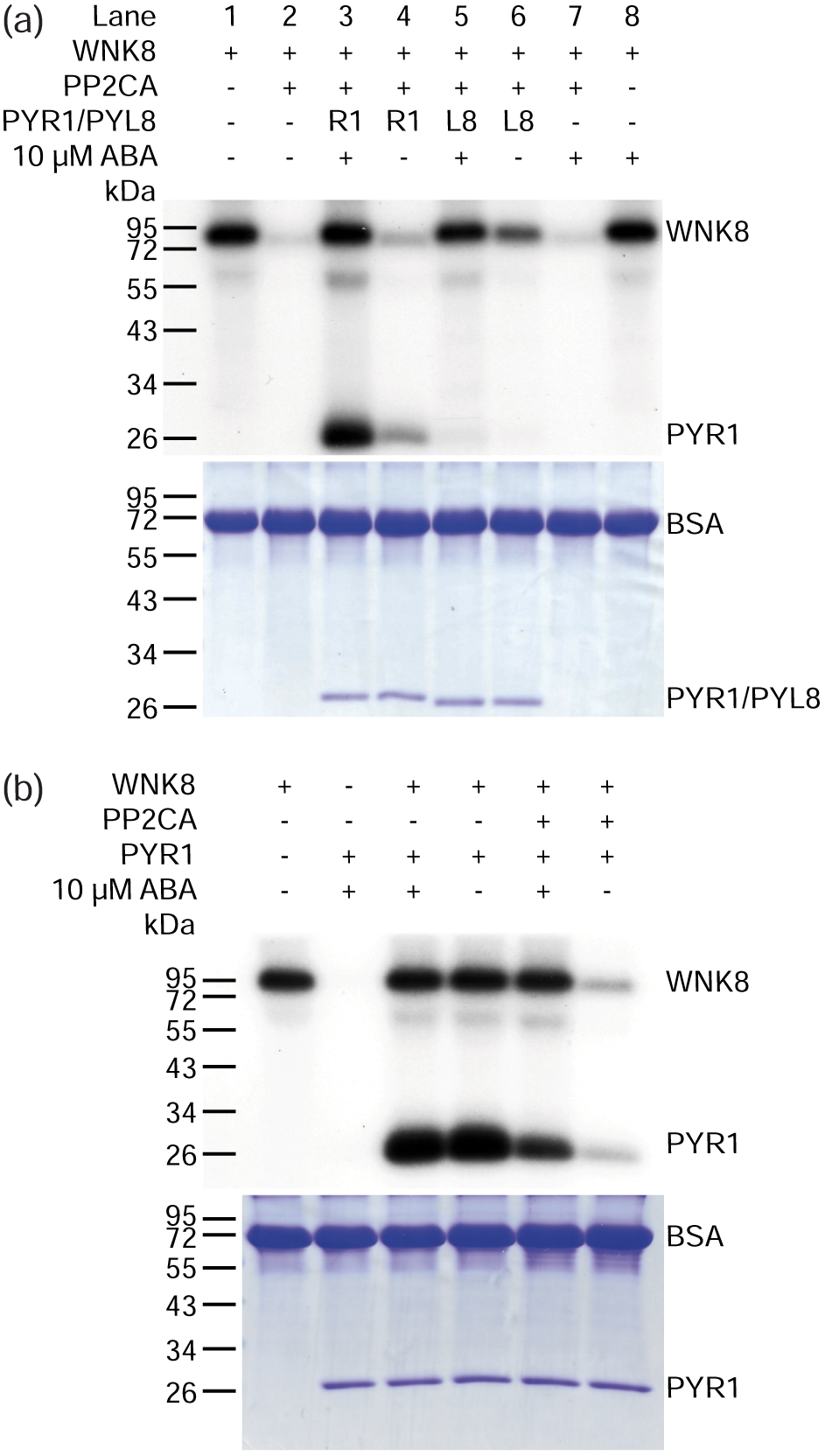
WNK8 phosphorylates itself and PYR1 and is negatively regulated by PP2CA *in vitro*. *In vitro* kinase assays of indicated protein and ABA mixtures (R1, PYR1; L8, PYL8). Top images, autoradiograph of the phosphorylated proteins. Bottom images, coomassie stained SDS-gels. Note that BSA was used as stabilizer for protein integrity. Kinase assays were performed for 30 min at 30°C and PP2CA incubations with PYR1, PYL8 and ABA were carried out on ice 20 min prior to WNK8 and ATP addition.

Consistent with previous reports (Hong-Hermesdorf et al., 2006), WNK8 alone exhibited auto-phosphorylation activity, independent of ABA (Figure 6a, lanes 1 and 8). However, in the presence of PP2CA, auto-phosphorylation of WNK8 was greatly reduced, indicating that PP2CA either inhibited WNK8 activity, or actively de-phosphorylated WNK8 (Figure 6a, lanes 2 and 7). The inhibition of WNK8 auto-phosphorylation by PP2CA was fully restored when 10 μM ABA and PYR1 or PYL8 were additionally added to the kinase assays (Figure 6a, lanes 3 and 5). In the absence of ABA, PYL8 had a minor effect on PP2CA inhibition (Figure 6a, lane 6), consistent with its ability to inhibit PP2CA also in the absence of ABA, when present in an increased molar ratio (Hao et al., 2011). Interestingly, inhibition of PP2CA by ABA and PYR1 enabled the phosphorylation of PYR1 by WNK8 (Figure 6a, lane 3). Under similar conditions, inhibition of PP2CA by PYL8 did not result in detectable PYL8 phosphorylation by WNK8 (Figure 6a, lane 5), indicating some degree of WNK8 target specificity. To confirm and strengthen our findings that WNK8 phosphorylates PYR1, we performed additional control experiments (Figure 6b). The data indicated that WNK8 did not require the presence of PP2CA and/or ABA for its ability to phosphorylate PYR1.

To obtain further details about WNK8 and PYR1 phosphorylation sites, we performed similar kinase assays in the absence of radio-labelled ATP and analyzed the phosphorylated proteins by mass spectrometry (Table S1 and Table S2). After purification of StrepII-WNK8 from wheat germ extracts, WNK8 was phosphorylated at several residues including the StrepII-tag (Table S1). Addition of ATP and subsequent mass spectrometric analyses identified WNK8 Thr244, Ser245, Ser251, Ser336 and Ser492 as potential WNK8 auto-phosphorylation sites (Table S1, highlighted in bold). Thr338 may represent an additional auto-phosphorylation site. However, mass spectrometric analyses resulted in a rather low localization probability, rendering this site ambiguous (Table S1). Interestingly, addition of PP2CA resulted in de-phosphorylation of most WNK8 phosphorylation sites (Table S1). WNK8 phosphorylated PYR1 at Ser85, Thr106, Ser109 and Thr162, and these sites were not de-phosphorylated by PP2CA (Table S2, highlighted in bold). Interestingly, PYR1_S85_ is located in the Pro-Cap, required to capture ABA in the bound conformation, and PYR1_S109_ and PYR1_T162_ are adjacent to residues critical for ABA binding (Nishimura et al., 2009). These data imply that PYR1 phosphorylation by WNK8 might affect ABA binding of PYR1 and thereby affecting its probability to inhibit PP2Cs.

## DISCUSSION

### *wnk8-1* most likely functions as a dominant allele

Based on mutant phenotypes plant WNK-type protein kinases have been implicated in the regulation of flowering time, circadian rhythm, glucose sensing and abiotic stress tolerance (Wang et al., 2008; Tsuchiya and Eulgem, 2010; Urano et al., 2012; Zhang et al., 2013; Fu et al., 2014). Earlier studies on *WNK9* overexpression and T-DNA insertion alleles pointed to its role as a positive regulator of ABA signaling (Xie et al., 2014). Our investigations of two *WNK8* T-DNA insertion alleles (Figure 1; Figure 4; Figure S3) and ABA response analyses in Arabidopsis protoplasts (Figure 5) revealed that WNK8 functions as a negative regulator of ABA and glucose responses. The opposing effects of the two alleles *wnk8-1* (SALK_024887; Wang et al., 2008) and *wnk8-4* (SALK_103318; *wnk8-1* in Urano et al., 2012) could be explained by a dominant, gain-of-function phenotype of *wnk8-1,* that harbours the T-DNA insertion in the C-terminal autoinhibitory domain, but still expresses the kinase domain. In contrast, the *wnk8-4* allele harbours the T-DNA insertion in the kinase domain and therefore most likely represents a typical loss-of-function allele (Figure 1a,b). The presence of hypermorphic protein kinase T-DNA insertion alleles is not unusual and has been reported recently also for the receptor-like kinase THESEUS1 (Merz et al., 2017). Although *wnk8-1* has been characterized regarding flowering time (Wang et al., 2008) and glucose responses (Urano et al., 2015), its dominant behaviour and gain of function nature have not been noted. Our data suggest that the reduced ABA-sensitivity of *wnk8-1* (Figure 1f,h) is probably due to expression of a constitutively active unregulated kinase lacking the C-terminal autoinhibitory domain. In other context in which the C-terminal domain of WNK8 is required, *wnk8-1* could indeed behave as a recessive, loss-of-function allele. In future studies it will thus be important to compare the phenotypes of both gain- and loss-of-function alleles. The opposing roles in ABA responses of WNK8 (this study) and WNK9 (Xie et al., 2014) is still puzzling. Our results point to a direct interaction of WNK8 with core ABA signaling components (Figure 2; Figure 3; Figure 6; discussed below). Thus it will be interesting to identify the direct link between WNK9 and the ABA signaling pathway.

### The role of WNK8 and PP2CA in glucose responses

Earlier studies identified WNK8 as a component of the core heterotrimeric G-protein interactome (Klopffleisch et al., 2011). Other components that were identified in this interactome screen were the scaffold proteins Receptor of Activated C Kinase1 (RACK1) and the Regulator of G-protein Signalling 1 (RGS1) (Klopffleisch et al., 2011). Both, RACK1 and RGS1 are involved in glucose responses and have been identified as substrates of WNK8 (Urano et al., 2012, 2015). In response to glucose, RGS1 is phosphorylated, most likely by WNK8 and other related WNK-type protein kinases (WNK1 and WNK10) that leads to the endocytosis of RGS1 from the plasma membrane and G-protein activation (Urano et al., 2012). RACK1 is a scaffold protein that interacts with both RGS1 and WNK8 (Klopffleisch et al., 2011). Besides its role as a negative regulator of ABA responses (Guo et al., 2009), the main proposed function of RACK1 is the regulation of ribosome biogenesis and protein translation (Guo et al., 2011; Islas-Flores et al., 2015). Phosphorylation of RACK1A by WNK8 most likely affects the protein stability of RACK1A and a RACK1A phospho-dead version was able to rescue the glucose hypersensitive phenotype of the *rack1a-2* mutant, while the phospho-mimetic RACK1A version did not (Urano et al., 2015). It was concluded that RACK1A acts downstream of WNK8 as the *rack1a-2/wnk8-1* double mutant exhibited a similar glucose hypersensitivity than the *rack1a-2* single mutant, while the *wnk8-1* allele displayed glucose hyposensitivity (Urano et al., 2015). In our experimental conditions *wnk8-1* was only slightly hyposensitive to 6% glucose when compared to Col-0 wild type, but was epistatic to *pp2ca-1* with the *wnk8-1/pp2ca-1* double mutant displaying a strong glucose hyposensitivity compared to the *pp2ca-1* single mutant (Figure S3). On the other hand, the *wnk8-4* allele exhibited a hypersensitive response to 6% glucose, similar to *pp2ca-1* (Figure S3). Glucose and ABA responses are connected on several levels (Finkelstein and Gibson, 2002; Figure 7) and our results point to an important role of PP2CA as a negative regulator of glucose responses (Figure S3). PP2CA could act on the glucose response pathway through the inhibition of WNK8 (Figure 6), but also via inhibition of the protein kinase SnRK1.1 (Rodrigues et al., 2013). SnRK1 sensed the energy status of the cell (Baena-González and Hanson, 2017) and functions through phosphorylation of the transcription factor bZIP63 (Mair et al., 2015). Interestingly, the group A PP2C AIP1/HAI2 is also involved in glucose responses, but opposite to *pp2ca-1*, *aip1* mutants exhibited a hyposensitive response to glucose (Lim et al., 2012).

**Figure 7.**
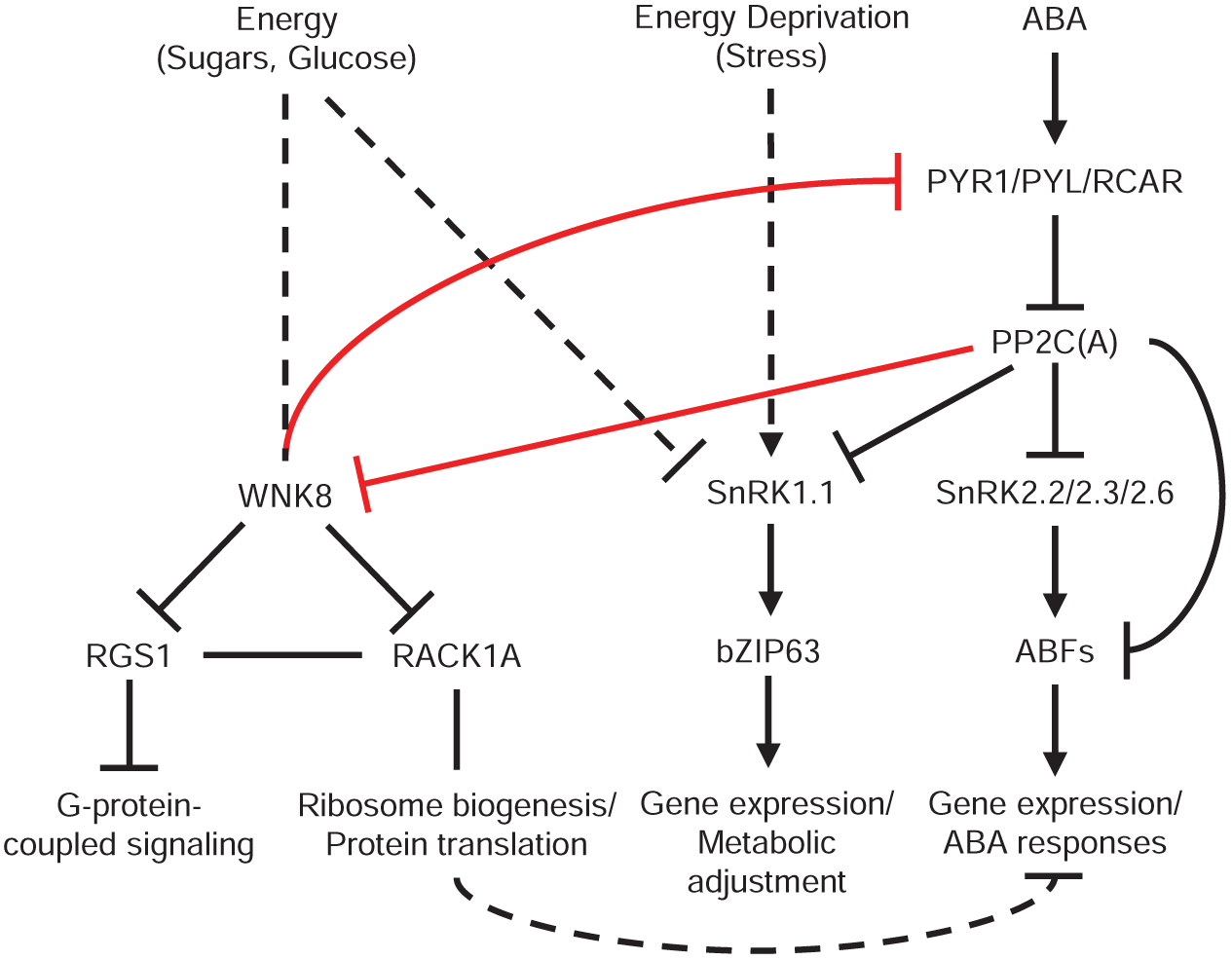
Hypothetical model of WNK8 function. (left) Known roles of WNK8 in the sugar sensing response. (middle) SnRK1 functions in sensing the energy status of a cell and is regulated by PP2Cs, including PP2CA. (right) The core ABA signaling pathway. Our identified connections between WNK8 and the ABA signaling pathway are indicated in red. Detailed description of the model is provided in the Discussion. Arrowhead, activation; perpendicular arrowhead, inhibition; line, unknown regulation; dashed line, indirect regulation.

### WNK8-mediated ABA responses are directly linked to interactions with core ABA signaling components

The core function of group A PP2Cs is the de-phosphorylation and negative regulation of SnRK2-type protein kinases (Cutler et al., 2010; Raghavendra et al., 2010; Hubbard et al., 2010). Without doubt SnRK2s are core players in ABA-signaling, but their interaction with PP2Cs is not exclusive. Members of the calcineurin B–like (CBL)–interacting protein kinase (CIPK)-family also interact with PP2Cs (Ohta et al., 2003) and share phosphorylation substrates with SnRK2s (Edel and Kudla, 2016). It has been also reported that CIPK23 can be directly de-phosphorylated by the PP2C ABI2 (Léran et al., 2015). The receptor-like kinase FERONIA is involved in various signaling pathways and functions as a negative regulator of ABA responses (Chen et al., 2016). Interestingly, although its negative role in ABA responses, FER is negatively regulated by ABI2 (Chen et al., 2016). Here, we report on similar findings regarding the protein kinase WNK8. Our data link the role of WNK8 as being a negative regulator of ABA responses (Figure 1; Figure 5) to its physical interaction with the core ABA signaling components PP2CA and PYR1 (Figure 2; Figure 3; Figure 6). We report here not only that inactivation of WNK8 by PP2CA is released by ABA in a receptor-dependent manner, but also that WNK8 is able to phosphorylate PYR1 *in vitro* (Figure 6). Several of the PYR1 residues that we have determined to be phosphorylated by WNK8 are in close proximity to the ABA-binding site (Table S2; Nishimura et al., 2009; Santiago et al., 2009a) and phosphorylation could thus affect the ABA-affinity of PYR1. To our knowledge phosphorylation of ABA-receptors has so far not been reported and future studies will be required to rigorously test this hypothesis.

A hypothetical model of WNK8 function could be that at high ABA concentrations, WNK8 can be activated and phosphorylates PYR1. PYR1 phosphorylation might negatively affect its ABA affinity or its ability to interact with PP2Cs. A reduced activity of PYR1 could release PP2CA from its inhibition by PYR1, thereby enabling the inhibition of WNK8 by PP2CA (Figure 7). In summary, we found that WNK8 functions as a negative regulator of ABA responses during young seedling development, and its role in ABA signaling was linked to its direct interaction with PP2CA and PYR1.

## EXPERIMENTAL PROCEDURES

### Plant materials and genotyping

Plant material used in this work is listed in Table S3. Genotyping was performed after isolation of genomic DNA from three-week-old plants using the CTAB method (Stacey and Isaak, 1994) and after RNA isolation from five-day-old seedlings using the RNeasy Plant Mini Kit (Qiagen). cDNA was reverse-transcribed from 2 μg of total RNA using MMLV-RTase (Fermentas) and oligo dT primers. Genomic PCRs and RT-PCRs were performed using oligo nucleotides listed in Table S4. Standard PCR reactions were performed using 29 PCR cycles for RT-PCRs and 35 cycles for genomic PCRs. T-DNA insertion sites were confirmed by sequencing of genomic PCR products.

### Plasmids

Plasmids (Table S5) were generated using oligo nucleotides listed in Table S4 and using standard cloning methods or the GreenGate system (Lampropoulos et al., 2013).

### Plant growth and seed germination assays

Seeds were surface sterilized in 70% EtOH for 10 min, washed three times in 100% EtOH, dried and sowed onto 0.5 MS media (Duchefa), 5 mM MES/KOH pH 5.8 solidified with 0.8% phytoagar (Duchefa). Seeds were stratified on the agar media for at least three days and seedlings were grown in a growth room (16 h day/8 h night, 22°C, 65% relative humidity, Philips Green Power LED deep red/blue 120 LO lamps, photon fluence rate 100 μmol m^−2^ s^−1^). Six-day-old seedlings were transferred to soil for seed propagation and genotyping.

For seed germination assays seeds were collected from single genotyped plants or from crossed siliques that have been grown in parallel. Seeds were sieved through a 400 μm mesh and debris smaller than 200 μm were removed. Blinded samples of 49 seeds were sown in quadruplicate onto 0.5 MS, 5 mM MES/KOH pH 5.8, 0.8% phytoagar media supplemented with or without 1% sucrose, indicated concentrations of ABA (TCI) or EtOH (control solution for ABA) and indicated glucose (Sigma-Aldrich) concentrations. After three days of stratification, plants were grown in the growth room and seed germination and cotyledon expansion or cotyledon expansion and greening was counted every day for 7-14 days. Data were normalized to the seed count (means ± SEM, n = 3-4), and graphs were generated using OriginPro.

For the rosette dry-down experiment plants were grown in a growth chamber (Conviron CMP 6010, 16 h day/8 h night, 22°C, 65% relative humidity, photon fluence rate 205 μmol m^−2^ s^−1^) after transfer to soil. Whole rosettes of three-week-old plants were cut-off, transferred to a weight boat and weighted every 15 min for 3 h and at time-points 6 and 9 h. Fresh weight was normalized to the initial weight at t = 0 min. Graphs were generated using OriginPro.

### *Promoter::GUS* analyses

*Promoter::GUS* analysis was performed on transgenic plants transformed with pCB308-WNK8pGUS (Table S5) using a promoter fragment of −1868 bp to +9 bp from the *WNK8* start codon. GUS-staining solution (50 mM NaPi, 0.1% Triton X-100, 20 mM X-Gluc and 2 mM K_3_FeCN_6_ and K_4_FeCN_6_) was vacuum infiltrated into whole seedlings using a standard water jet pump. Samples were stained at 37°C in darkness for several hours and stopped by sample incubation in 70% EtOH.

### Transient expression in *N. benthamiana* and microscopic analyses

Plant expression vectors for sub-cellular localization, BiFC and FRET-FLIM analyses, listed in Table S5, were transformed into the *Agrobacterium tumefaciens* strain ASE containing the pSOUP plasmid and infiltrated into leaves of four-to-five week old *Nicotiana benthamiana* plants grown in a greenhouse. Infiltration was conducted using standard protocols (Waadt et al., 2014). BiFC analyses (Waadt et al., 2008) were performed at 3 dpi while sub-cellular localization and FRET-FLIM analyses were conducted at 4 dpi. For FRET-FLIM analyses, *Agrobacteria* were infiltrated at an OD_600_ of 1 and WNK8 fusion proteins were expressed under control of a p*UBQ10* driven beta-estradiol-inducible promoter (Schlücking et al., 2013). Induction was carried out at 3 dpi by painting the abaxial leaf side with 100 μM beta-estradiol (Sigma).

Microscopic analyses were performed at an inverted Leica TCS SP5 II equipped with HCX PL APO CS 20x 0.7 IMM and HCX PL APO lambda blue 63x 1.2 WATER objectives and guided by the LAS AF software version 2.7.4.10329.

For BiFC analyses, images were acquired using identical microscope settings (1024×1024 pixels resolution, 514 nm excitation and 525-560 nm emission (BiFC); 561 nm excitation and 580-640 nm emission (mCherry)). Image analyses and processing was performed using Fiji (Schindelin et al., 2012). For the visualization of protein complex localizations, brightness and contrast of representative YFP (BiFC) images was optimized. To enable the visualization of BiFC emission differences images were color-coded and calibrated to a royal look up table using Fiji (Schindelin et al., 2012). For graphical analyses, BiFC emission was quantified from nuclei (n ≥ 20) and from entire acquired images (n = 20). Box plots were generated using OriginPro.

For FRET-FLIM analyses, the donor (GFP) and acceptor (mCherry) fluorescence intensity images were acquired with 512×512 pixels resolution, GFP and mCherry were excited simultaneously with 488 and 561 nm and emission was detected between 495-540 nm (GFP) and 600-640 nm (mCherry). Fluorescence lifetime data of GFP were acquired using a pulsed laser (50 ps pulse length), a time-correlated single photon counting board (PicoHarp 300, Picoquant) and an integrated FLIM-photomultiplier tube (R7400U, Hamamatsu). GFP was excited at 470 nm with a pulse frequency of 20 MHz and the emitted photons were collected between 490-550 nm. Lifetime images were acquired at 256×256 pixels resolution with 100 repetitions per measurement. Fluorescence decay curves were calculated from time resolved fluorescence intensity images and fitted to a bi-exponential decay model using the Symphotime software v5.2.4.0 (Picoquant).

### ABA signaling analyses in Arabidopsis protoplasts

Preparation of Arabidopsis protoplasts and ABA signaling analysis was performed as described (Moes et al., 2008; Ma et al., 2009). Information about plasmids used for protoplast transformations is provided in Table S5. The protoplast suspensions were incubated at 25°C in the absence or presence of 10 μM exogenous ABA, and reporter expression was determined after 16 to 18 h. Assays were carried out in triplicates per data point and presented data derive from three biological replicates.

### Protein purifications and *in vitro* kinase assays

Plasmids used for protein expression are described in Table S5. His-tagged PYR1 and PYL8 were expressed in M15 pREP4 and BL21(DE3) cells. Cells were grown in liquid LB medium at 37°C until they reached OD_600_ of 0.5 and proteins were expressed for 4 h after induction with 0.75 mM (PYR1) or 1 mM (PYL8) of IPTG. Protein purification was performed using Protino^®^ Ni-TED 2000 Packed Columns (Macherey & Nagel) according to the manufacturer’s protocol. PYR1-6xHis was re-buffered into 100 mM Tris-Cl pH 7.6, 100 mM NaCl, 0.3 mM MnCl_2_, 4 mM DTT and 6xHis-PYL8 was re-buffered into 50 mM Tris-Cl 7.5, 0.1% Tween20, 10 mM beta-mercaptoethanol using Amicon Ultra-0.5 mL Centrifugal Filters (Millipore). StrepII-tagged PP2CA and WNK8 proteins were expressed and purified according to Hashimoto et al. (2012). For storage, the purified proteins were supplemented with 1 mg mL^−1^ BSA.

*In vitro* kinase assays were performed according to Hashimoto et al. (2012). Purified proteins (200 ng WNK8, 400 ng PP2CA, and same amounts of PYR1 and PYL8) were mixed as indicated in the experiment and volume was adjusted to 16 μL with water. PP2CA and PYR1 or PYL8 were incubated on ice for 20 min with or without 10 μM (+/-) ABA (Sigma) before addition of WNK8. After incubation, samples were supplemented with 4 μL 6x reaction buffer (30 mM MnSO_4_, 3 mM CaCl_2_, 12 mM DTT) and kinase reaction was started by addition of 4 μL of 6x ATP mixture (60 μM ATP magnesium salt, 1 μCi μl^−1^ [gamma-^32^P] ATP) (Hartmann Analytic). 30 min after incubation at 30°C reactions were stopped by 1 μl addition of 0.5 M EDTA and 6 μl of 5x Lämmli buffer (250 mM Tris-Cl pH 6.8, 10% SDS, 0.05% bromophenol blue, 10% beta-mercaptoethanol, 40% glycerol). After 5 min incubation at 95°C, samples were separated by SDS-PAGE and SDS-gels were stained using coomassie brilliant blue, scanned for documentation of protein amounts and exposed to x-ray films at room temperature to capture phosphorylated proteins.

### Mass spectrometric analyses

Description of the mass spectrometric analyses in provided in Methods S1.

## ACCESSION NUMBERS

*ABI1* (AT4G26080); *ACTIN2* (AT3G18780); *HSP18.2* (AT5G59720); *PP2CA* (AT3G11410); *PYR1* (AT4G17870); *PYL8* (AT5G53160); *RD29B* (AT5G52300); *UBQ10* (AT4G05320); *WNK8* (AT5G41990).

## ACQNOWLEDGEMENTS

We thank Pedro Rodriguez for the pET28a-PYL8 plasmid. This work was funded by grants from the Deutsche Forschungsgemeinschaft to MH, EG, JK and KS (FOR964).

## AUTHOR CONTRIBUTIONS

KS, JK, EG and MH conceived the project, EG, AHH, RW, YL, KH, MK and MS performed the experiments and analyzed the data, RW and KS wrote the manuscript and all other authors revised the manuscript.

## SUPPORTING INFORMATION

**Methods S1** Mass spectrometric analyses.

**Table S1.** WNK8 phosphorylation sites identified by mass spectrometry.

**Table S2.** PYR1 phosphorylation sites identified by mass spectrometry.

**Table S3.** Transgenic lines used in this work.

**Table S4.** Oligonucleotides used in this work.

**Table S5.** Plasmids generated in this work.

## FIGURE LEGENDS

**Figure S1.**
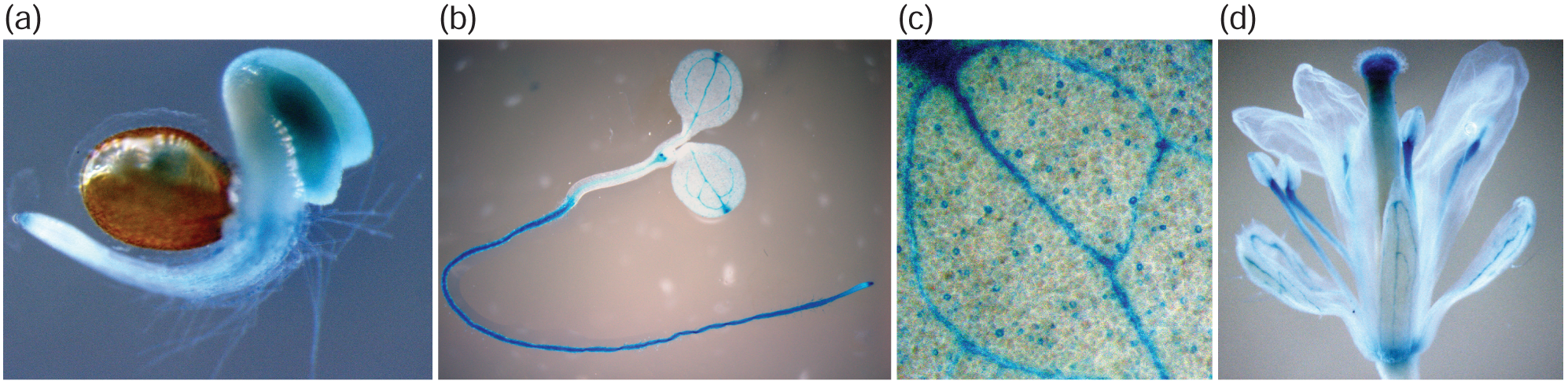
*Promoter::GUS* analysis of the WNK8 gene. *Promoter::GUS* analysis using a −1868 bp to +9 bp promoter fragment of the WNK8 gene. (a) two-day-old seedling, (b) five-day-old seedling, (c) closeup on a cotyledon of a five-day-old seedling, and (d) flower.

**Figure S2.**
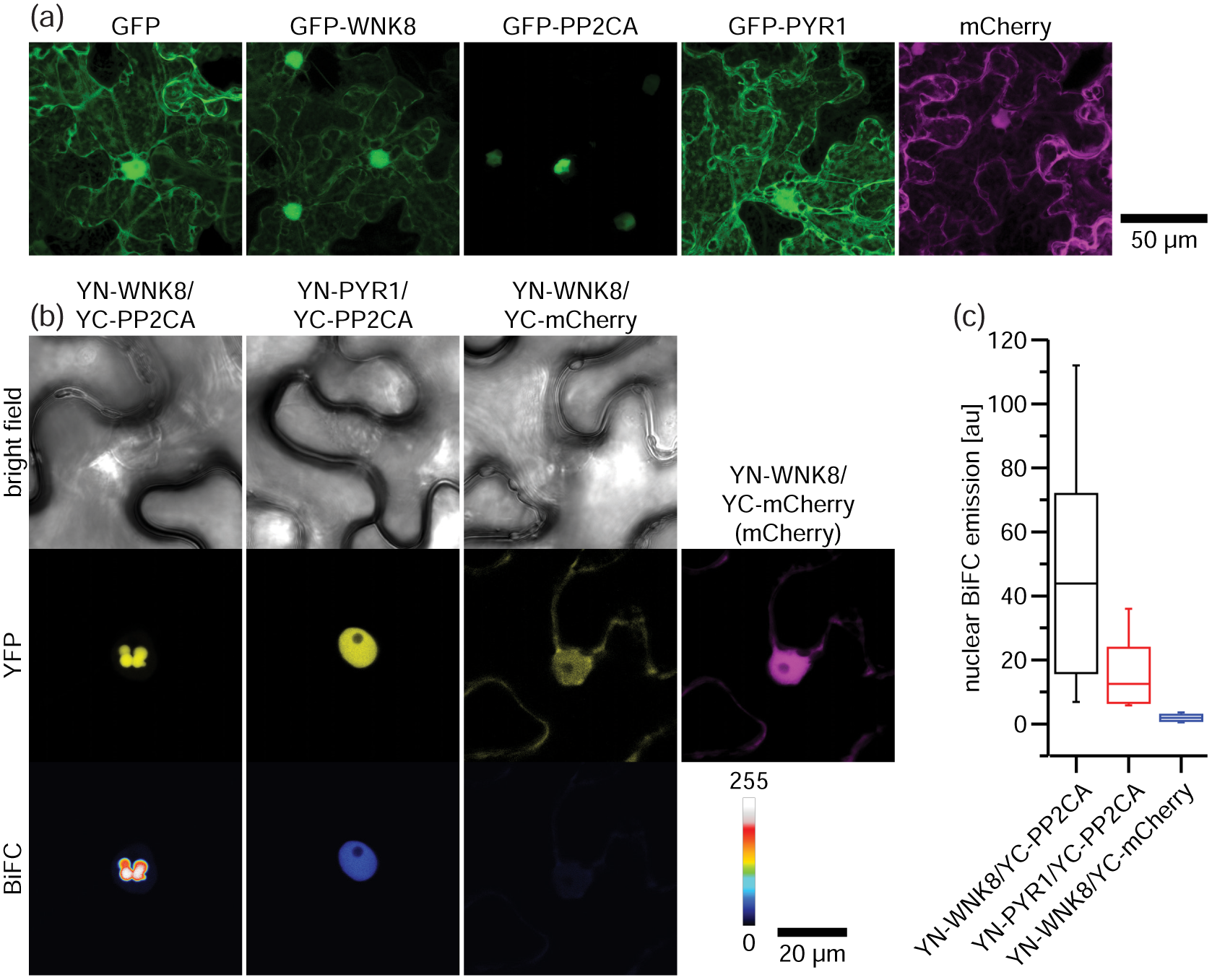
Additional subcellular localizations and BiFC analyses. (a) Subcellular localization of indicated constructs after transient expression in N. benthamiana leaf epidermis cells. Shown are representative maximum z-projection images of 32 z-frames. (b) From top to bottom, representative bright field, subcellular localization (YFP) and comparative BiFC emission (BiFC) of indicated protein complex combinations after transient expression in N. benthamiana leaf epidermis cells. BiFC emissions were calibrated to the adjacent color bar. Expression of YC-mCherry is indicated in magenta. (c) Quantification of nuclear BiFC emissions from indicated protein complex combinations. Boxes, median ± SD; whiskers, minimum and maximum values (n ≥ 20). See also Figure 2 for related BiFC analyses.

**Figure S3.**
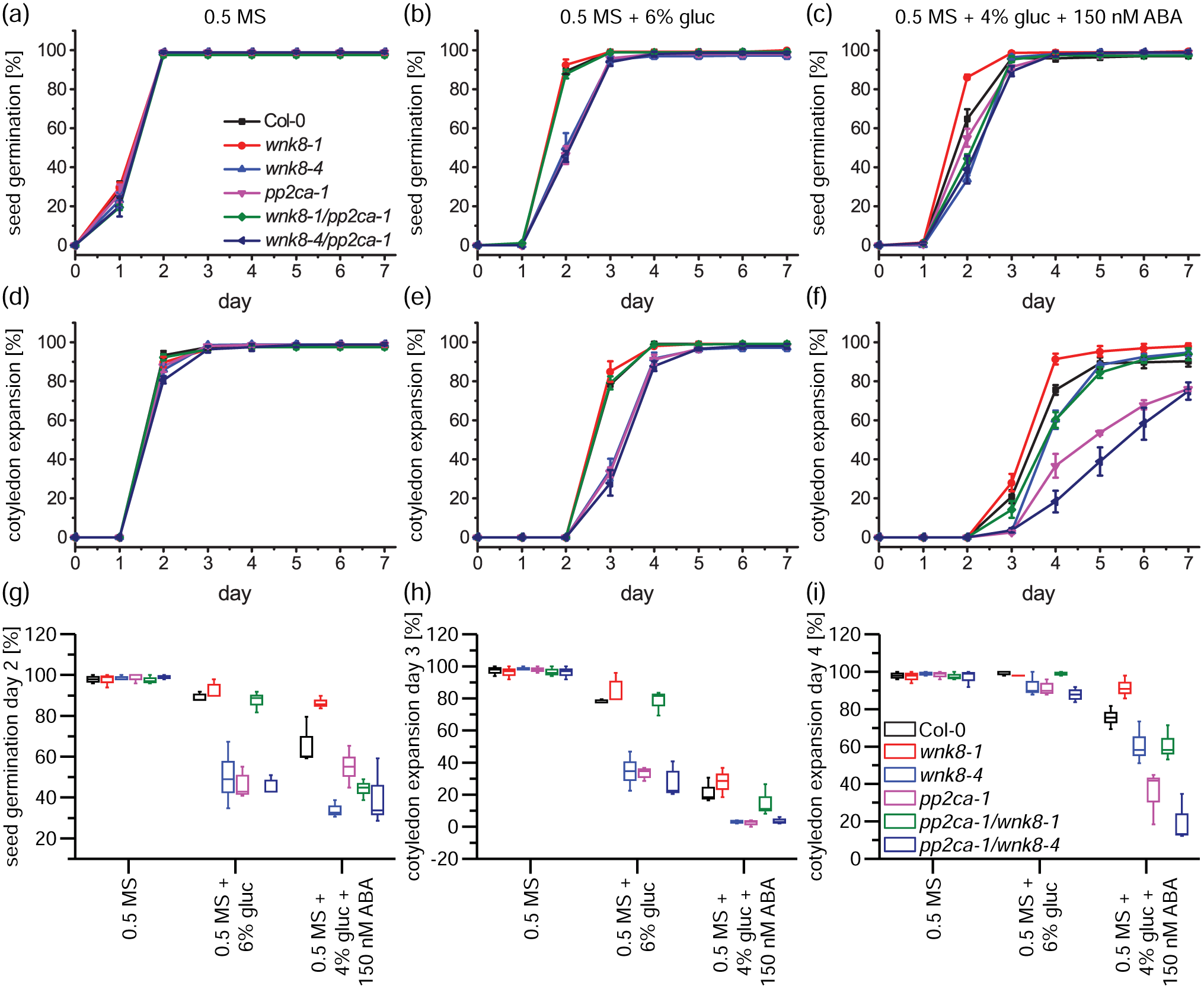
WNK8 and PP2CA mediate glucose and ABA responses during young seedling development. (a-c) Time course of seed germination and (d-f) cotyledon expansion of indicated genotypes when grown on 0.5 MS media supplemented with (a,d) 0.015% EtOH as solvent control, (b,e) 6% glucose (gluc) and 0.015% EtOH, or 4% glucose and 150 nM ABA. (g-i) Box plots from data in (a-f) taken at indicated time points. Boxes, median ± SD; whiskers, minimum and maximum values (n = 3-4). See also Figure 4 for related experiments.

